# An essential experimental control for functional connectivity mapping with optogenetics

**DOI:** 10.1101/2022.05.26.493610

**Authors:** David Tadres, Hiroshi M. Shiozaki, Ibrahim Tastekin, David L. Stern, Matthieu Louis

## Abstract

To establish functional connectivity between two candidate neurons that might form a circuit element, a common approach is to activate an optogenetic tool such as Chrimson in the candidate pre-synaptic neuron and monitor fluorescence of the calcium-sensitive indicator GCaMP in a candidate post-synaptic neuron. While performing such experiments, we found that low levels of leaky Chrimson expression can lead to strong artifactual GCaMP signals in presumptive postsynaptic neurons even when Chrimson is not intentionally expressed in any particular neurons. Withholding all-*trans* retinal, the chromophore required as a co-factor for Chrimson response to light, eliminates GCaMP signal but does not provide an experimental control for leaky Chrimson expression. Leaky Chrimson expression appears to be an inherent feature of current Chrimson transgenes, since artifactual connectivity was detected with Chrimson transgenes integrated into three different genomic locations (two insertions tested in larvae; a third insertion tested in the adult fly). These false-positive signals may complicate the interpretation of functional connectivity experiments. We illustrate how a no-Gal4 negative control improves interpretability of functional connectivity assays. We also propose a simple but effective procedure to identify experimental conditions that minimize potentially incorrect interpretations caused by leaky Chrimson expression.

## Introduction

A primary goal of neuroscience is to provide a mechanistic understanding of behavior at the level of circuits of connected neurons. In recent years, the combination of precise anatomical reconstruction of neuronal circuits (Dorkenwald et al., 2022; Lillvis et al., 2021; Meinertzhagen, 2018; Scheffer et al., 2020; Schlegel et al., 2017) and the ability to genetically target sparse subsets of neurons has provided new opportunities to decipher neural computation (Borst and Helmstaedter, 2015; Eschbach et al., 2021; Randel et al., 2015; Takemura et al., 2013; Wang et al., 2021). In *Drosophila melanogaster*, large collections of transgenic flies have been created to reproducibly label or target effector proteins to small subsets, and sometimes individual, neurons in the fly brain (Hayashi et al., 2002; Jenett et al., 2012; Li et al., 2014; Tirian and Dickson, 2017). Genetically-encoded effectors are typically expressed in subsets of cells using binary expression systems such as UAS-Gal4 (Brand and Perrimon, 1993), lexAop-lexA (Lai and Lee, 2006) and QUAS-QF (Potter et al., 2010). To test whether candidate neurons are involved in a given behavior, standard methodologies of neurogenetics include constitutive loss-of-function assays with a modified version of the tetanus toxin (TNT) or the inwardly-rectifying potassium (Kir) channel, and acute gain-of-function assays with tools from thermogenetics and optogenetics (Simpson, 2009; Simpson and Looger, 2018).

Neuronal activity is often assayed by monitoring fluorescence of genetically-encoded indicators such as the calcium sensor GCaMP (Chen et al., 2013), voltage sensors such as ASAP (Chamberland et al., 2017) and neurotransmitter sensors (Borden et al., 2020; Jing et al., 2020). More recently, reconstruction of neuronal connectivity in serial section electron microscopy (Takemura et al., 2013) and FIB-SEM (Scheffer et al., 2020) has provided putative connectomes of parts of the adult fly (Scheffer et al., 2020; Schlegel et al., 2017; Takemura et al., 2013) and the larva (Schneider-Mizell et al., 2016). From connectomic data, neural circuit diagrams can be constructed to formulate hypotheses about functional connectivity between neurons (Eschbach and Zlatic, 2020; Gowda et al., 2021). For example, we recently used an EM connectome of the larval nervous system to identify the downstream partner of a command like descending neuron that leads to stopping behavior in fruit fly larvae (Tastekin et al., 2018).

To test hypotheses about functional connectivity, separate binary expression systems are used to express optogenetic gain-of-function tools (for example, Chrimson) in one set of cells and a neuronal activity reporter (for example, GCaMP) in putative downstream neurons. If the optogenetic activation of the presynaptic neuron leads to reproducible signals in the post-synaptic neuron, it is often concluded that the two neurons are connected either directly or indirectly, thereby forming a circuit element. A common experimental control for these experiments is to withhold the chromophore all-*trans* retinal (ATR), which is required for Chrimson activity, from the diet of flies (see Materials and methods for more details). If putative downstream neurons in flies fed ATR show increased activity upon optogenetic activation of upstream neurons and do not show increased activity when ATR is withheld (ATR^-^), then the increased activity of putative downstream neurons is dependent on optogenetic activation (Carreira-Rosario et al., 2018; Takagi et al., 2017; Yoshino et al., 2017). This control is adequate to test for innate light responses. However, it does not assess the possibility that the activation is caused by basal or “leaky” expression from the UAS-Chrimson transgene (expression in the absence of a Gal4 driver), which could lead to false detection of functional connectivity. While the presence of basal expression has been reported for some UAS transgenes (Markstein et al., 2008), it is unclear if basal Chrimson expression is sufficient to generate detectable GCaMP signal in downstream neurons in the absence of a Gal4 driver, especially when neurons are exposed to relatively high activating light levels.

We found that UAS-Chrimson transgenes inserted into several genomic locations exhibit leaky expression that can cause significant calcium responses in functional connectivity experiments. These responses are not detected in flies lacking ATR. Therefore, the calcium responses observed in ATR^-^ flies represent false-positive signals that can generate misleading conclusions about the connectivity of pairs of pre- and post-synaptic neuron candidates. To avoid conflating calcium transients originating from leaky expression with true positive signals, it is crucial to test closely-matched genetic controls that do not contain a Gal4 driver.

## Results

### Leaky expression can lead to ambiguous and false evidence for circuit connectivity

Using connectomics data from electron microscopy, we identified neurons upstream and downstream of the PDM descending neuron (PDM-DN). One downstream candidate was SEZ-DN1 (Tastekin et al., 2018). To establish that SEZ-DN1 is functionally downstream of PDM-DN, we expressed the optogenetic tool UAS-CsChrimson::tdTomato (hereafter abbreviated as UAS-Chrimson) in PDM-DN and the fluorescent reporter GCaMP in the SEZ-DN1 neuron. We used a parental control lacking the Gal4 driver with and without the addition of ATR as negative controls (Figure 1A). Surprisingly, we found a significant increase in calcium signal, measured as relative change in fluorescence with respect to baseline (ΔF/F), in the ATR^+^ negative control but not in the ATR^-^ negative control (Figure 1B and 1C). The signal found in the ATR^+^ negative controls is thus a false positive, since the UAS-Chrimson transgene was not intentionally expressed in any neuron. False-positive signals were found when the preparation was stimulated with a range of red light intensities overlapping those often used in functional connectivity experiments (Figure 1E and 1F).

**Figure 1:**
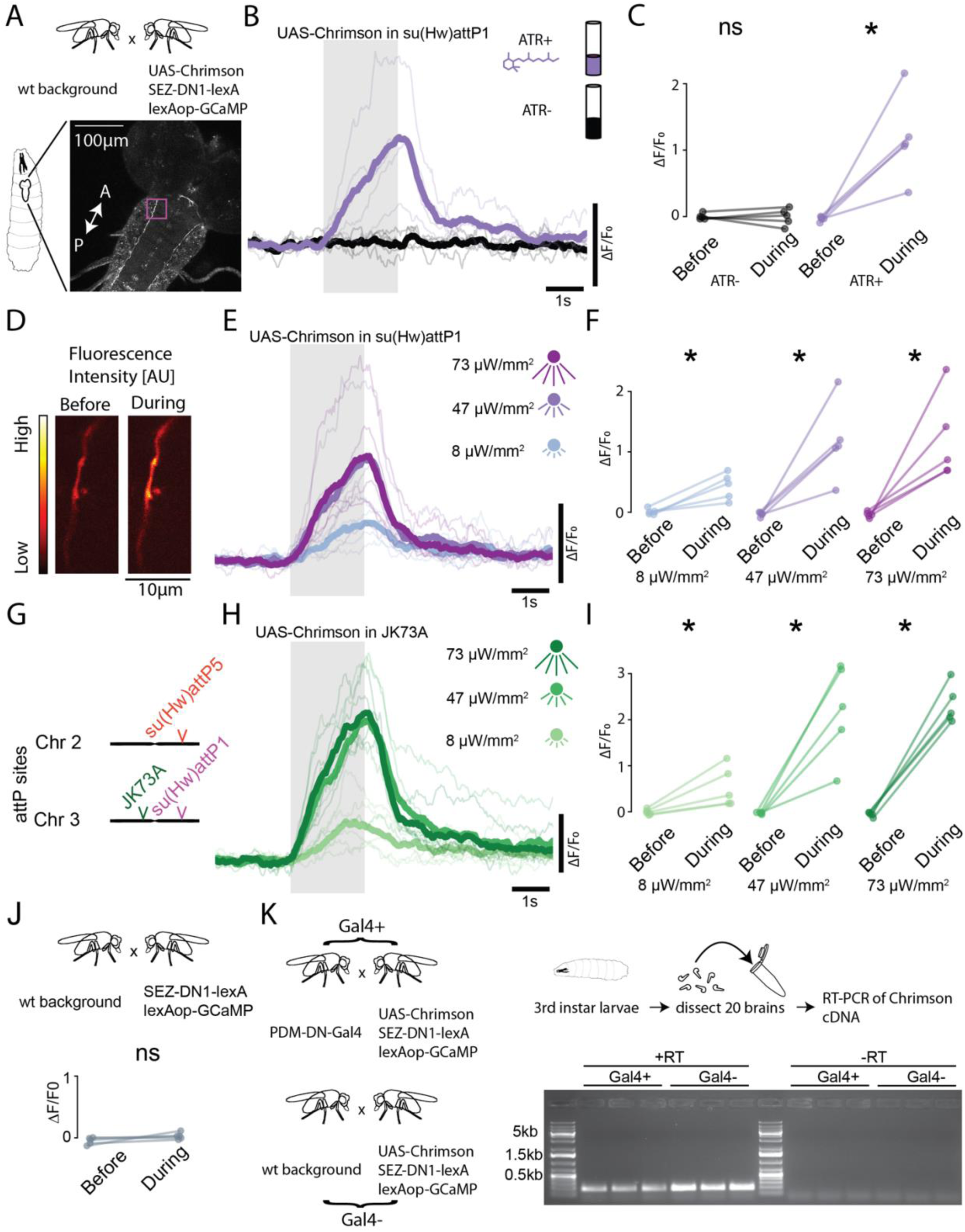
Leaky expression of UAS-Chrimson is the cause of false-positive calcium signal in a descending pathway of the *Drosophila* larva. **(A)** Calcium imaging was performed on larvae of no-Gal4 parental control flies. **(B)** Calcium transient of larvae grown on food with all-*trans* retinal (magenta) and on food devoid of all-*trans* retinal (black). Grey shade indicates 590 nm stimulus with power of 47 uW/mm2. **(C)** Larvae with the UAS-Chrimson transgene in su(Hw)attP1 have a significant increase in ΔF/F upon stimulation (magenta). Larva grown on food devoid of all-trans retinal have no detectable calcium transients. The star indicates a statistically significant difference (n=5 animals, paired t-test, p<0.05 upon Holm-Bonferroni correction) **(D)** Example image of pixel intensity before (left) and during (right) stimulation. **(E)** Calcium transients of larvae grown on food with all-trans retinal at 3 different stimulus light intensities, 21 uW/mm2 (light blue), 47 uW/mm2 (lilac) and 72 uW/mm2 (purple). **(F)** The calcium transients for all three stimulus light intensities are significantly higher during stimulation compared to before. (n=5 animals, paired t-test, p<0.05 upon Holm-Bonferroni correction). **(G)** In this study, UAS-Chrimson transgenes were considered in three different landing sites: su(Hw)attP1 (magenta), JK73A (green) and su(Hw)attP5 (red, Figure 2). **(H)** Calcium transient of larvae with UAS-Chrimson in JK73A landing site grown on food with all-trans retinal at 3 different stimulus light intensities, 8 uW/mm2 (light green), 47 uW/mm2 (green) and 72 uW/mm2 (dark green). **(I)** The calcium transients for all three stimulus light intensities shown in panel H are significantly higher during stimulation compared to before stimulation (paired t-test, p<0.05 upon Holm-Bonferroni correction). **(J)** In larvae without any UAS-Chrimson transgene no significant change is observed during stimulation (paired t-test, p>0.05) **(K)** For each biological replicate, 20 brains of 3^rd^ instar were dissected and the mRNA was extracted. The mRNA was used to create cDNA. The expected band size for cDNA of Chrimson is 155bp. Both larvae with and without Gal4 produce cDNA of Chrimson. The no-reverse transcriptase control (-RT) shows that the primers used were specific to cDNA of Chrimson. The raw calcium traces underlying this figure are presented in Supplementary Fig. 1. In panels 1B, 1E and 1H, the thick lines represent the means of the calcium signals computed over the medians of individual trials per experiment (thin lines).

To determine whether this false-positive signal resulted from an endogenous ATR-dependent light-sensitive protein, we tested larvae devoid of the UAS-Chrimson transgene but fed ATR (Figure 1J). We found no significant calcium transients. Thus, the false-positive signals in SEZ-DN1 require the presence of the UAS-Chrimson transgene, and may reflect leaky expression of Chrimson.

Transgene expression is influenced by the location of its insertion in the genome (Pfeiffer et al., 2010). To test whether potentially leaky Chrimson expression was specific to transgene insertions at su(Hw)attP1, we repeated our analysis with a UAS-Chrimson transgene inserted into a different landing site that supports robust expression, JK73A (Figure 1G) (Knapp et al., 2015). We observed false-positive signals over a range of light intensities for flies carrying a UAS-Chrimson transgene at JK73A (Figure 1H and 1I). These results indicate that false-positive signals can result from putative leaky expression of the UAS-Chrimson transgene inserted at multiple genomic locations.

To test whether leaky expression arises from the UAS-Chrimson transgene, we conducted RT-PCR on brains of larvae with and without a Gal4 transgene (Gal4^+^ and Gal4^-^, respectively) to drive the expression of UAS-Chrimson. In three independent replicates, cDNA of Chrimson was detected irrespective of the presence of the Gal4 driver (Figure 1K). The results of the RT-PCR corroborate the hypothesis that the false-positive signal is due to leaky expression of the UAS-Chrimson transgene undirected by any Gal4 driver.

### False-positive signals are found at the larval and the adult stages

Given the observation of leaky expression of Chrimson at the larval stage, we next asked whether the same artifact can be observed at the adult stage. We expressed GCaMP in a subset of Kenyon cells that innvervate the gamma (γ) lobe of the mushroom bodies, ventral accessory calyx, and peduncle (γd Kenyon cells) (Aso et al., 2014). These Kenyon cells convey visual information and underlie visual learning (Vogt et al., 2016). In flies lacking any Gal4 driver, we observed significant calcium responses in the γ lobe of the mushroom bodies of adult flies (Figure 2A and 2D). As we observed for larvae (Figure 1), significant calcium transients (ΔF/F) were found in control flies lacking the Gal4 driver when raised with ATR, but not in absence of ATR (Figure 2B and 2C). We observed a significant increase in calcium signal for light intensities ranging from 72 to 1151 uW/mm^2^ (Figure 2E and 2F). We also imaged the calyx and peduncle of the mushroom body (Supplementary Fig. 2A) and observed a significant increase in calcium signal when stimulating with 1151 uW/mm^2^ (Supplementary Fig. 2B and 2C). To test if the false-positive signal originated from an endogenous ATR-dependent light-sensitive protein, we imaged from adult flies devoid of the UAS-Chrimson transgene, but raised on ATR. In these conditions, we found no significant calcium transients (Supplementary Fig. 2D), showing that the calcium transients observed in the γ lobe of the mushroom bodies are Chrimson-dependent. Together, these results show that leaky expression of a UAS-Chrimson transgene can lead to false-positive signals in a widely studied neuronal population in the adult brain.

**Figure 2:**
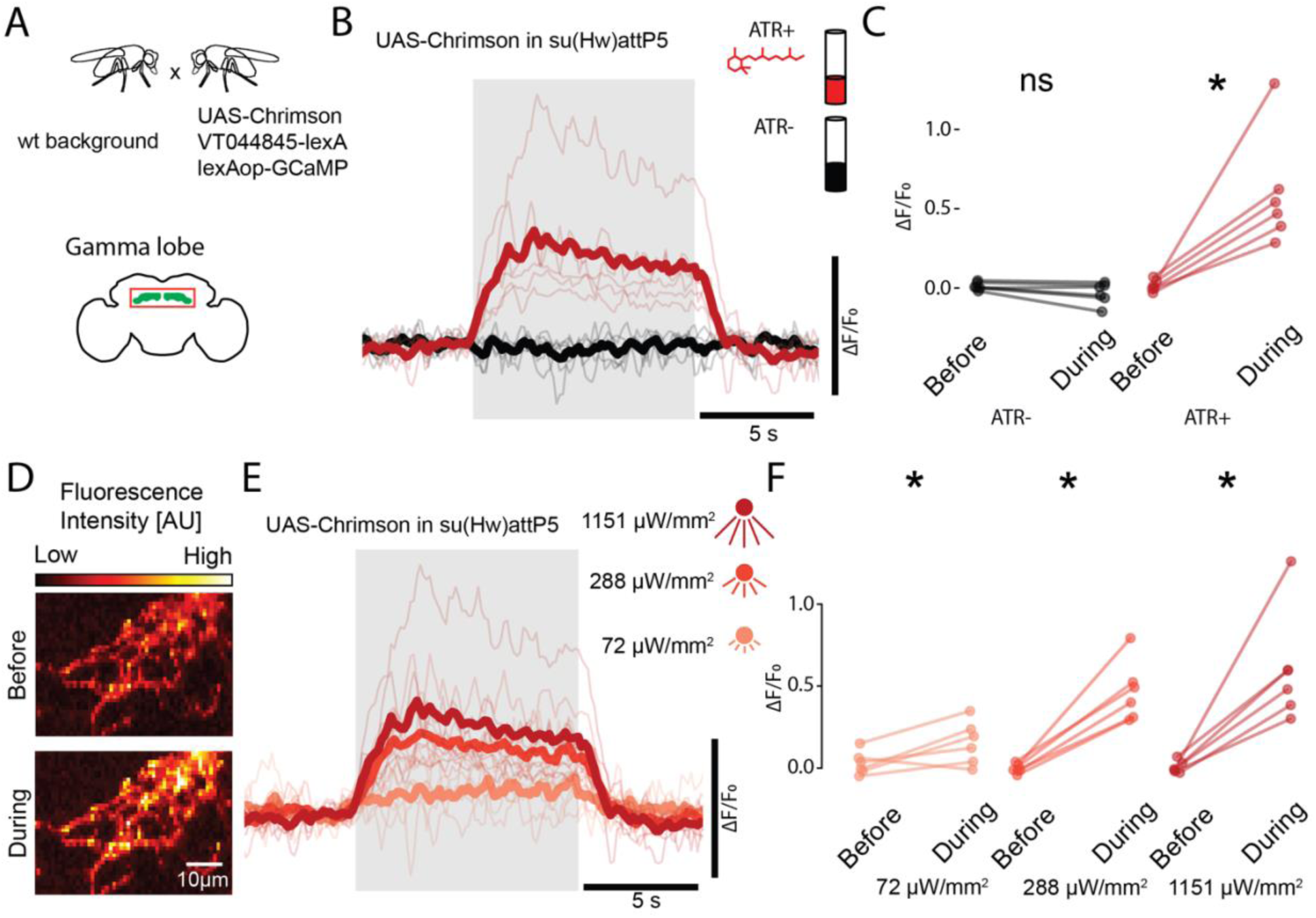
Leaky expression of UAS-Chrimson is the cause of false-positive calcium signal in the mushroom body of the Drosophila adult. **(A)** Calcium imaging was performed in the γ lobe of adult flies that lack a Gal4 driver. **(B)** Calcium transients of larvae grown on food with all-trans retinal (red) and on food devoid of all-trans retinal (black). Grey shade indicates 660 nm stimulus with power of 1151 uW/mm2. **(C)** Flies with the UAS-Chrimson transgene in su(Hw)attP5 have a significant increase in ΔF/F upon stimulation when fed with ATR (red). Flies grown on food devoid of ATR have no detectable calcium transients (n=6 animals, paired t-test, p<0.05 upon Holm-Bonferroni correction). **(D)** Example image of pixel intensity before (top) and during (bottom) stimulation in part of the γ lobe in one hemisphere. **(E)** Calcium transients of flies grown on food with ATR at 3 different stimulus light intensities, 72 uW/mm2 (light red), 288 uW/mm2 (red) and 1151 uW/mm2 (dark red). **(F)** The calcium transients for all three stimulus light intensities shown in panel F are significantly higher during stimulation compared to before stimulation (n=6 animals, paired t-test, p<0.05 upon Holm-Bonferroni correction). The raw calcium traces underlying this figure are presented in Supplementary Fig. 2E. In panels 2B and 2E, the thick lines represent the means of the calcium signals computed over the medians of individual trials per experiment (thin lines).

### Validating true positive signals in control experiments to rule out the effects of leaky expression

Although leaky expression of Chrimson can lead to false-positive calcium signals (Figures 1 and 2), these results do not invalidate this approach for functional connectivity. One control to establish connectivity is to test for a calcium response due to the leaky expression of Chrimson in flies lacking any Gal4 driver (Gal4^-^, ATR^+^) in parallel with the test condition with a Gal4 driver (Gal4^+^, ATR^+^). By comparing the response magnitude between the control and test conditions, one can estimate how much of the response reflects Chrimson expression driven by a Gal4 driver (Figure 3A-C).

**Figure 3:**
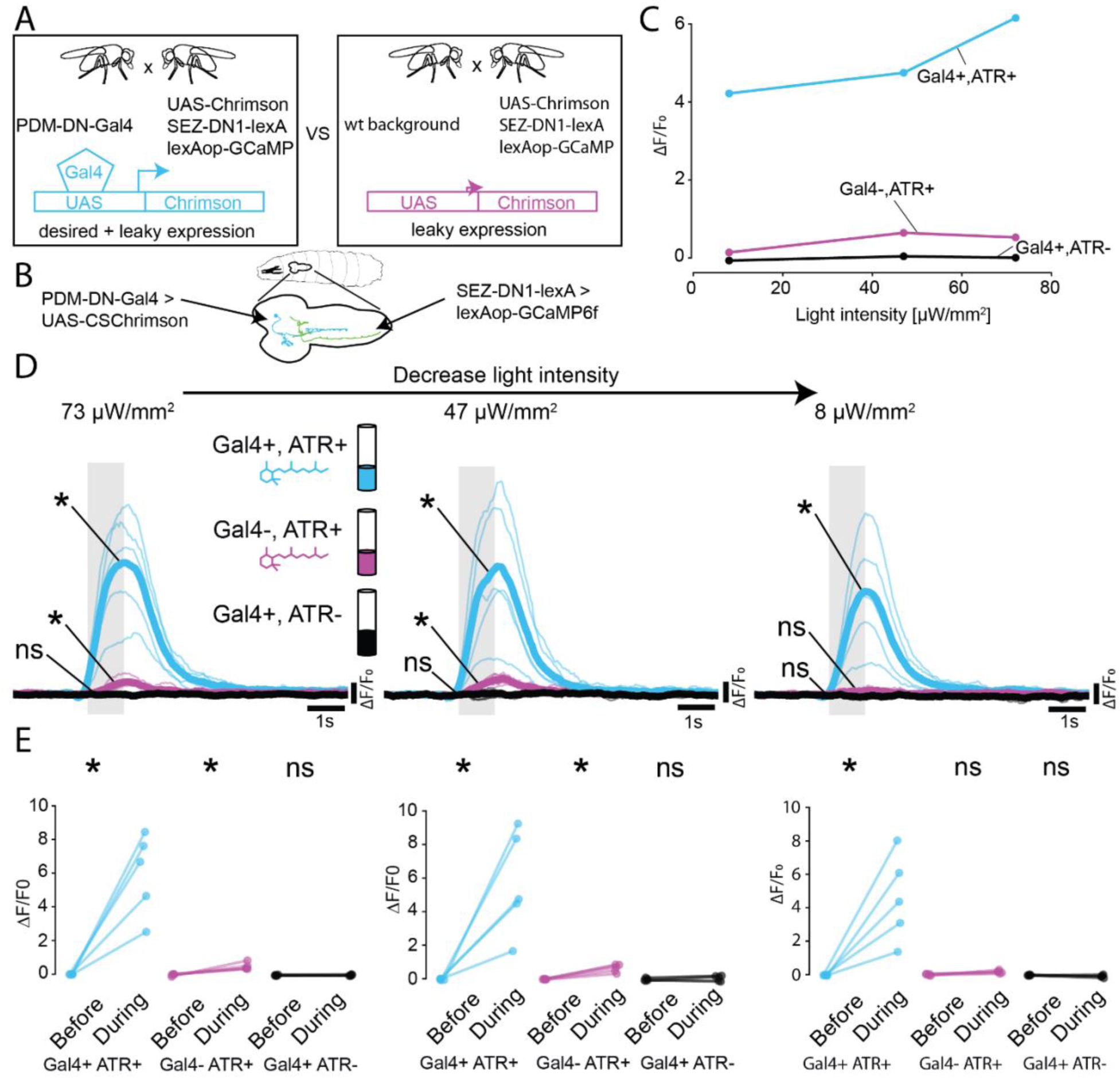
Testing a no-Gal4 parental control raised on retinal represents a robust strategy to rule out false-positive calcium signals due to leaky expression. **(A)** To maximize confidence in calcium signal measured in experiments aimed at establishing connectivity between pre- and post-synaptic partners, the experimenter can set up a no-Gal4 control (Gal4-,ATR+) in parallel to the test genotype (Gal4+,ATR+). This allows comparing signal arising from the leaky expression of Chrimson and the Gal4-driven expression to identify conditions of stimulation where the signal due to the leaky expression is negligible. **(B)** In this example, Chrimson is expressed in the PDM-DN neuron which has been shown to be upstream of SEZ-DN1 neuron (Tastekin et al., 2018). **(C)** Illustration of the dose response of the median calcium transient responses of the no-Gal4 (bottom) and Gal4 test (top) conditions over an order of magnitude of stimulus intensity for 2-second light stimulations. **(D)** At 73 uW/mm2 and at 47 uW/mm2 both the No-Gal4 control and Gal4 test conditions have a calcium transient different from zero. At 8 uW/mm2 the calcium transient of the Gal4 test condition is different from zero while the no-Gal4 control is not. **(E)** Comparisons of the peak calcium transients before and during the light stimulation (n=5 animals, paired t-test, p<0.05 upon Holm-Bonferroni correction). Light stimulation of a duration of 1 second and an intensity 8 uW/mm2 represents a trade-off where true-positive calcium signals are detected, and false-positive signals are avoided. The raw calcium traces underlying this figure are presented in Supplementary Fig. 3. In panel 3D, the thick lines represent the means of the calcium signals computed over the medians of individual trials per experiment (thin lines).

The effect of leaky UAS expression can be minimized by systematically searching for a light stimulation intensity in which signals associated with the Gal4 driver, but not false-positive signals, are observed. We illustrate the application of this procedure to detect connectivity between PDM-DN and SEZ-DN1 (Figure 3D and 3E). We tested a range of decreasing light intensitites from 73 to 8 uW/mm^2^ to identify a Chrimson stimulation regime where true signal can be differentiated from signal resulting from leaky expression. We found that activation with 8 uW/mm^2^ produced a significant response in the Gal4^+^, ATR^+^ flies, but not in Gal4^-^, ATR^+^ flies, suggesting that the GCaMP signal reflects connectivity between the PDM-DN neurons expressing Chrimson and the SEZ-DN1 neurons expressing GCaMP.

## Discussion

Recent advances in electron microscopy and big-data analysis have provided candidate wiring diagrams for whole brains or parts thereof in *Drosophila* at the larval and adult stages(Buhmann et al., 2021; Eichler et al., 2017; Hulse et al., 2021; Scheffer et al., 2020; Schneider-Mizell et al., 2016; Wanner et al., 2016; Zheng et al., 2018). After inferring that two neurons are potentially connected from electron microscopy data, the next logical step is to establish functional connectivity between the two neuron candidates (Franconville et al., 2018; Simpson and Looger, 2018). This can be done efficiently in *Drosophila* by expressing an optogenetic tool in the putative upstream neuron while monitoring the calcium activity of the putative downstream neuron. We found that candidate downstream neurons can generate significant calcium responses associated with false-positive signals in parental controls lacking any Gal4 driver but raised with retinal supplement (Figure 1).

While we focused on two examples, the technical problem we reported is likely to be widespread and common to many functional connectivity experiments. We linked the existence of false-positive signals to the basal — “leaky”— expression of the UAS-Chrimson transgene. We showed that false-positive signals occur both in larvae and adults in commonly studied cells such as descending neurons (Figures 1 and 3) and in the mushroom body (Figure 2). Consistent with the fact that multiple landing sites are susceptible to leaky expression of UAS transgenes (Markstein et al., 2008), we found evidence for leaky Chrimson expression from UAS transgenes inserted in three different genomic locations (Figure 1G). Moreover, the parameters of optogenetic stimulation that induced false-positive responses in our experiments were within the range used in published studies. For example, false-positive responses were observed at a light intensity of 8 uW/mm^2^ in the larva (Figure 1E) and 72 uW/mm^2^ in the adult (Figure 2F) where much stronger light intensities have been used previously (for instance, 680 uW/mm^2^ (Chen et al., 2021); 2170 uW/mm^2^ (Sen et al., 2017)). Given that retinal supplementation is required for Chrimson to function in flies (Klapoetke et al., 2014), it is not surprising that false-positive calcium signals were never observed in flies carrying both Gal4 and UAS-Chrimson transgenes in the absence of all-*trans* retinal (Figure 3C-E). Consequently, withholding retinal from flies is not a sufficient control for optogenetic-based connectivity experiments, since this treatment does not control for leaky transgene expression. Other controls should be included in optogenetic-based connectivity experiments.

To validate that calcium signals are caused by the optogenetic activation of an upstream candidate neuron, we propose the following control experiment. First, a line without Gal4 and raised on retinal (Gal4^-^, ATR^+^) should be assayed under identical conditions as the genetically-matched line containing Gal4 (Gal4^+^, ATR^+^). Second, it is recommended to use stimulation conditions where only the experimental genotype (Gal4^+^, ATR^+^) yields a significant increase in calcium signal (ΔF/F). Such conditions can be found by gradually changing the intensity of the light stimulus (Figure 3D and 3E). In addition, false-positive responses can also be reduced by decreasing the duration of the light stimulus as illustrated by the comparison between the signal elicited by a 2-second and a 1-second presentation of light at 8 uW/mm^2^ (Figure 1E and Figure 3E, respectively). Thus, identifying the stimulation conditions where only the experimental condition (Gal4^+^, ATR^+^) yields a significant signal represents an effective safeguard against false-positive artifacts in the assessment of connectivity between candidate neurons forming a circuit.

The cautious approach required to interpret the results of functional manipulations based on Chrimson activation applies also to loss-of-function and gain-of-function behavior experiments. This is particularly relevant to screens with a behavioral readout where the leaky expression of the effector can produce unexcepted changes in behavior. For example, Scholz et al. (2000) observed higher than baseline ethanol sensitivity in UAS-TNT flies in the absence of Gal4. Tang et al. (2017) found increased temperature preference in UAS-TrpA1-RNAi flies in the absence of Gal4. Our results emphasize the importance of performing control experiments with the UAS effector line alone in loss-of-function experiments involving TNT (Keller et al., 2002; Scholz et al., 2000), Kir2.1 or RNAi (Qiao et al., 2018; Tang et al., 2017), as well as in gain-of-function experiments using TrpA1 to produce thermogenetic activation (Simpson, 2009). In contrast to these experiments, where the experimenter has little or no control over the strength of the effector, the efficacy of the activation (conductance) of Chrimson can be modulated by the light stimulation protocol. This distinctive property enables any experimenter to mitigate false-positive signals resulting from the leaky expression of Chrimson.

Mapping functional circuits is a prerequisite for understanding the mechanisms of neural computation. As the number of driver lines for expressing effector proteins in specific cell types in *Drosophila* has increased, functional connectivity mapping with optogenetics has become increasingly popular. Here, we demonstrate that the leaky expression of the optogenetic actuator can compromise the integrity of such circuit mapping. Fortunately, a simple experiment with flies carrying no Gal4 driver and a careful comparison of GCaMP activity over a range of activation intensities provides a suitable control strategy that can be easily integrated in future efforts to map functional connectivity.

## Materials and methods

### Fly rearing

It is usually necessary to complement food with all-*trans* retinal when conducting optogenetic experiments in *Drosophila* (Klapoetke et al., 2014). However, cornmeal contains β-carotene and zeaxanthin (De Oliveira and Rodriguez-Amaya, 2007) which are precursors of all-*trans* retinal (Oberhauser et al., 2008). To completely eliminate all-*trans*-retinal in our negative controls (“ATR^-^”), we prepared fly food using ‘Nutri-Fly GF’ (Genesee Scientific, #66-115) which is devoid of cornmeal. All-*trans* retinal (R2500-500MG, Sigma Aldrich) was added to the ‘Nutri-Fly GF’ fly food to yield a final concentration of 0.5 mM (“ATR^+^”).

### Flies used in this study

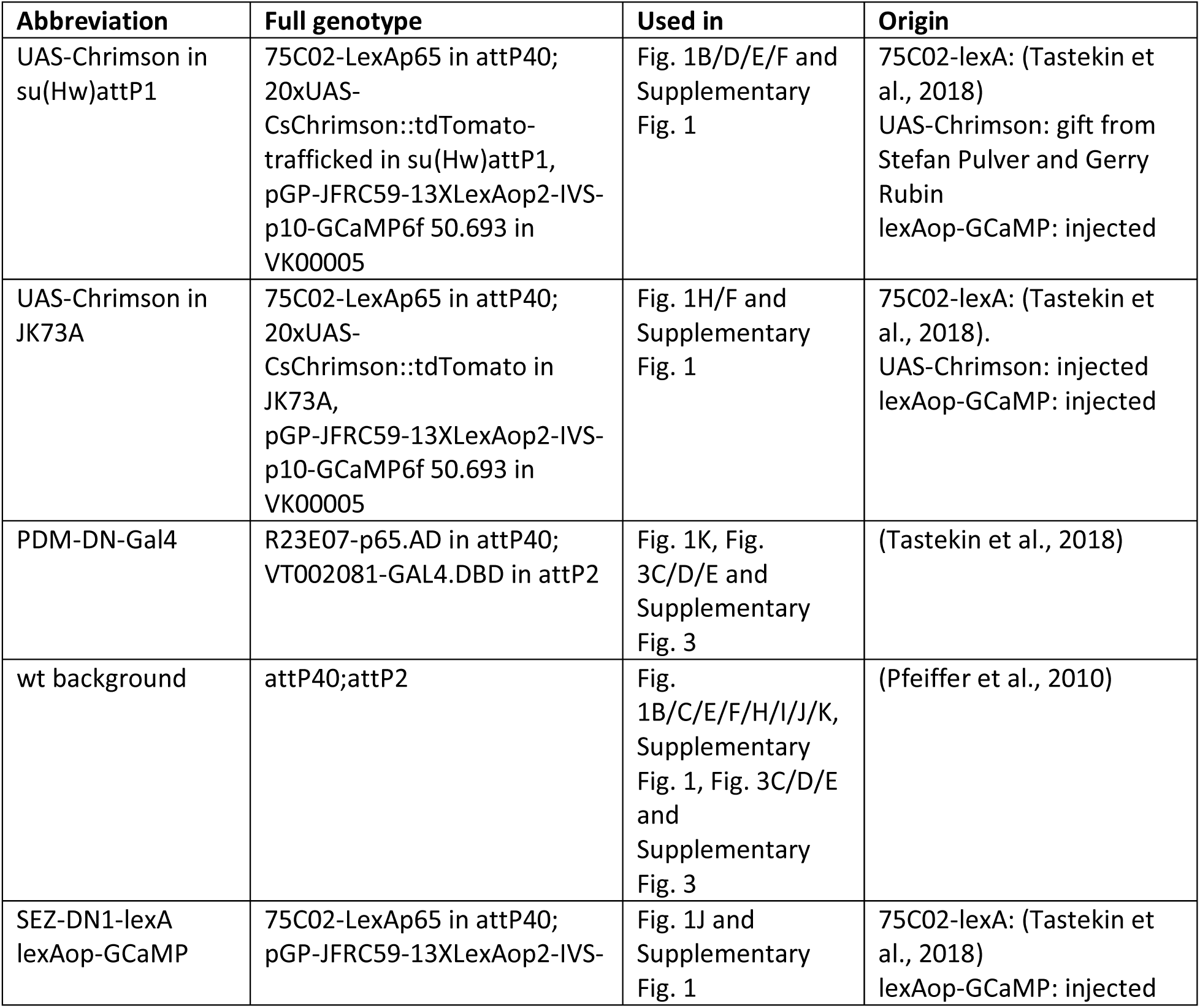
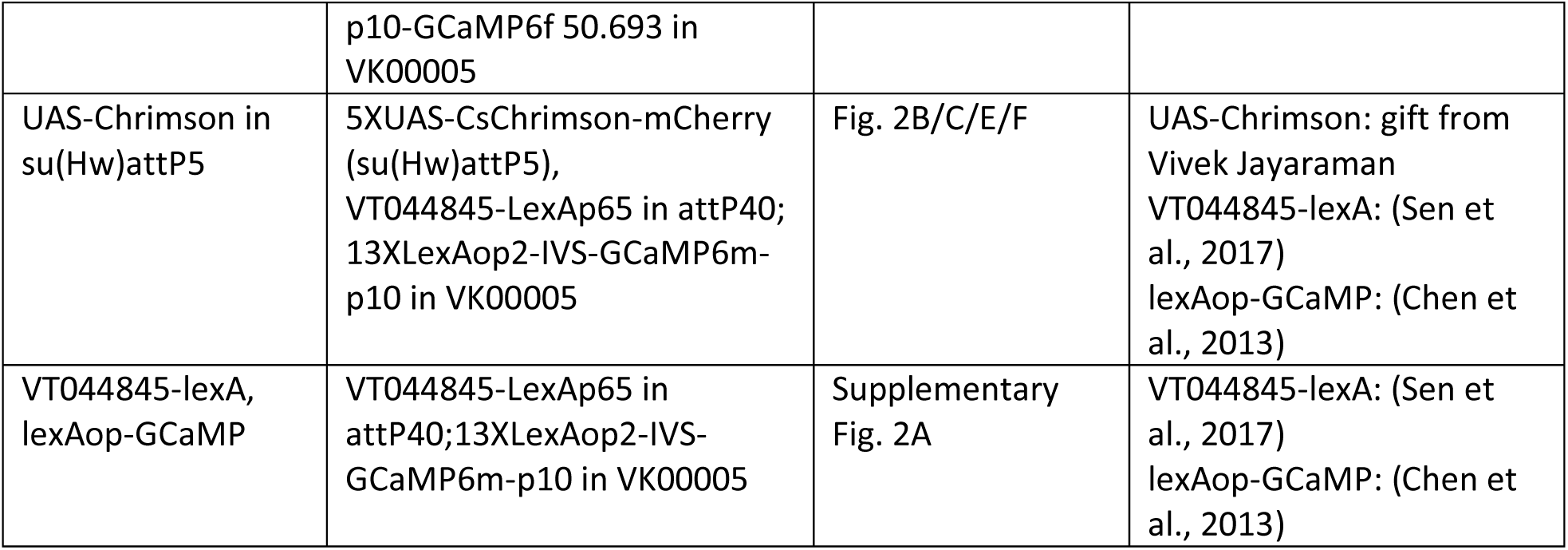

| Abbreviation | Full genotype | Used in | Origin |
| --- | --- | --- | --- |
| UAS-Chrimson in su(Hw)attP1 | 75C02-LexAp65 in attP40; 20xUAS-CsChrimson::tdTomato-trafficked in su(Hw)attP1, pGP-JFRC59-13XLexAop2-IVS-p10-GCaMP6f 50.693 in VK00005 | Fig. 1B/D/E/F and Supplementary Fig. 1 | 75C02-lexA: (Tastekin et al., 2018)<br>UAS-Chrimson: gift from Stefan Pulver and Gerry Rubin<br>lexAop-GCaMP: injected |
| UAS-Chrimson in JK73A | 75C02-LexAp65 in attP40; 20xUAS-CsChrimson::tdTomato in JK73A, pGP-JFRC59-13XLexAop2-IVS-p10-GCaMP6f 50.693 in VK00005 | Fig. 1H/F and Supplementary Fig. 1 | 75C02-lexA: (Tastekin et al., 2018).<br>UAS-Chrimson: injected<br>lexAop-GCaMP: injected |
| PDM-DN-Gal4 | R23E07-p65.AD in attP40; VT002081-GAL4.DBD in attP2 | Fig. 1K, Fig. 3C/D/E and Supplementary Fig. 3 | (Tastekin et al., 2018) |
| wt background | attP40;attP2 | Fig. 1B/C/E/F/H/I/J/K, Supplementary Fig. 1, Fig. 3C/D/E and Supplementary Fig. 3 | (Pfeiffer et al., 2010) |
| SEZ-DN1-lexA<br>lexAop-GCaMP | 75C02-LexAp65 in attP40; pGP-JFRC59-13XLexAop2-IVS- p10-GCaMP6f 50.693 in VK00005 | Fig. 1J and Supplementary Fig. 1 | 75C02-lexA: (Tastekin et al., 2018)<br>lexAop-GCaMP: injected |
| UAS-Chrimson in su(Hw)attP5 | 5XUAS-CsChrimson-mCherry (su(Hw)attP5), VT044845-LexAp65 in attP40; 13XLexAop2-IVS-GCaMP6m-p10 in VK00005 | Fig. 2B/C/E/F | UAS-Chrimson: gift from Vivek Jayaraman<br>VT044845-lexA: (Sen et al., 2017)<br>lexAop-GCaMP: (Chen et al., 2013) |
| VT044845-lexA, lexAop-GCaMP | VT044845-LexAp65 in attP40;13XLexAop2-IVS-GCaMP6m-p10 in VK00005 | Supplementary Fig. 2A | VT044845-lexA: (Sen et al., 2017)<br>lexAop-GCaMP: (Chen et al., 2013) |

### Plasmids used in this study

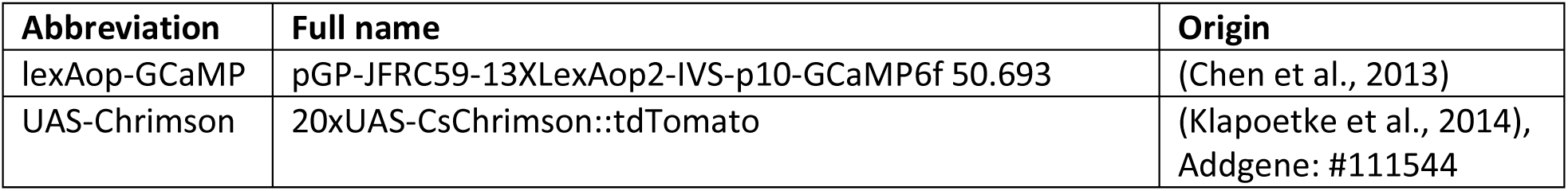

### cDNA preparation (larva)

For each biological replicate, 20 larval brains were dissected. RNA was extracted with *the ReliaPrep RNA Miniprep System* (Promega, Z6111) which includes a DNA digestion step with *DNAse I*. RNA amount and quality was measured using a *NanoDrop* (ThermoFisher). *SuperScript IV Reverse Transcriptase* Kit (ThermoFisher, 18090010) was used with 500 ng total RNA and oligoT primers to create cDNA. This was followed by RNA digestion step using *RNAse H*. In the -RT control, water was used instead of Reverse Transcriptase.

### RT-PCR (larva)

*Q5 High-Fidelity DNA Polymerase* (NEB, M0491L) was used with indicated primers. PCR was performed with annealing temperature of 65C and extension time 30 seconds with 30 cycles.

### List of Primers

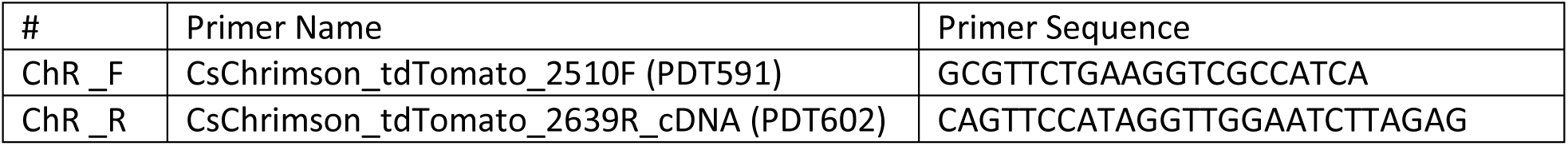

### Calcium imaging in the larva

The CNS of larvae at the 3rd instar developmental stage were dissected in ‘Pulver Saline’ (135 mM NaCl, 5 mM KCl, 4 mM MgCl2, 1.8 mM Trehalose, 12.6mM Sucrose, 2mM CaCl2) and placed dorsal side down on a lysine-coated coverslip. We used an Hyperscope two-photon scanning microscope (Scientifica, UK) with a MaiTai laser (Spectra Physics, CA, USA). For imaging, we used two-photon excitation wavelength of 920 nm. CsChrimson was activated with a LED (LCS-0590-03-22, Mightex, Canada) with peak wavelength at 590 nm and a 605/55 bandpass filter (Chroma, VT, US). To feed the CsChrimson excitation light into the light path, we used a 560LP dichroic (T560LPXRXT-UF1, Chroma, VT, USA). The power of the LED from the objective was measured with a PM100D Power meter with a S170C Power sensor (Thorlabs, NJ, USA). To estimate the power at the larval brain we divided the measured total LED power by the area of the field of view of the objective (XLPLN25XWMP2, Olympus, Japan).

For each trial, the neuron was observed for 20 seconds. Each trial was repeated 5 times. Data was analyzed using custom python scripts. Raw data was first filtered using a Savitzky-Golay filter (1-second window length, 2^nd^ order polynomial). The signal was calculated as described previously (Jia et al., 2011). Briefly, a ROI was chosen to contain a segment of the axonal branch. A second ROI away from the neuron was defined as the background. The background was subtracted from the ROI to correct for stimulus light bleed-through. Next, for each frame F_i_, signals were computed as the relative change in fluorescence intensity from the baseline: ΔF/F= (F_i_-F_0_)/F_0_. Baseline fluorescence F_0_ was defined as the mean pixel intensity of the 3 second time window preceding the optogenetic stimulus.

Preliminary experiments showed that many but not all preparations show calcium transients. For the larva (Figures 1 and 3), the experiments were performed using the following algorithm: each dissected brain was stimulated using the same 2-second 73 uW/mm2 step stimulus. If an obvious response was detected the whole dataset was acquired. If no obvious response was detected, the brain was discarded and another brain was dissected and tested. For the 6^th^ preparation in a row, imaging was performed irrespective of the presence of a response to initial 2-second step stimulus. This allowed us to efficiently scan through 30 samples to minimize the chances of missing false positives in unexpected conditions. The table below shows the number of brains dissected and recorded.

| Genotype | all-*trans*-retinal | Figure | Brains dissected | Data collected |
| --- | --- | --- | --- | --- |
| UAS-Chrimson in su(Hw)attP1, SEZ-DN1-lexA, lexAop-GCaMP | ATR<sup>+</sup> | Figure 1B-F<br>Figure 3C-E | 6 | 5 |
| UAS-Chrimson in su(Hw)attP1, SEZ-DN1-lexA, lexAop-GCaMP | ATR<sup>-</sup> | Figure 1B | 30 | 5 |
| UAS-Chrimson in JK73A, SEZ-DN1-lexA, lexAop-GCaMP | ATR<sup>+</sup> | Figure 1H and 1I | 7 | 5 |
| SEZ-DN1-lexA, lexAop-GCaMP | ATR<sup>+</sup> | Figure 1J | 30 | 5 |
| PDM-DN-Gal4, UAS-Chrimson in su(Hw)attP1, SEZ-DN1-lexA, lexAop-GCaMP | ATR<sup>+</sup> | Figure 3D-F | 7 | 5 |
| PDM-DN-Gal4, UAS-Chrimson in su(Hw)attP1, SEZ-DN1-lexA, lexAop-GCaMP | ATR<sup>-</sup> | Figure 3D-F | 30 | 5 |

### Calcium imaging in the adult

The CNS of 5-7 day old adult female flies were dissected in saline (103 mM NaCl, 3 mM KCl, 5 mM N-tris(hydroxymethyl) methyl-2aminoethane-sulfonic acid, 8 mM trehalose, 10 mM glucose, 26 mM NaHCO3, 1 mM NaH2PO4, 1.5 mM CaCl2, and 4 mM MgCl2; pH, 7.1-7.3; osmolarity, 270-275 mOsm). The saline was bubbled with carbogen (95% O2 and 5% CO2) prior to the dissection. For the recording from the γ lobe of the mushroom bodies, the sample was placed on a coverslip so that the anterior brain faced upward. For the recording from the calyx and peduncle of the mushroom bodies, the posterior brain faced upward. Calcium imaging was conducted with a two-photon microscope (Bergamo II, Thorlabs) equipped with a 16x objective lens (N16XLWD-PF, Nikon) and a pulsed laser tuned to 940 nm (InSight X3, Spectra-Physics). The laser power measured under the objective lens was below 25 mW. Samples were imaged at 10 z-planes covering 40 μm with a Piezo scanner (PFM450E, Thorlabs). Each z-plane was imaged every 140.8 ms (7.1 volumes per s). CsChrimson was activated with a 660-nm laser (S1FC660, Thorlabs) delivered through an optic fiber placed at 3 mm away from the sample (M125L01, Thorlabs). The power of the CsChrimson activation laser at the sample was calculated under the assumption that the power was uniformly distributed within the beam radius. The laser power varied from 72 to 2302 uW/mm2. For each trial, the laser at a fixed power was continuously applied to the sample for 10 s with the inter-trial interval of 20 s. The laser of the same power was applied for 6 trials.

Time series of stack images underwent rigid motion correction with NoRMCorre (Pnevmatikakis and Giovannucci, 2017). These images were averaged across z-planes for the following analyses. An ROI that contained the target neurons and a ROI for background were manually defined based on the images averaged across the entire recording. ΔF/F was calculated with baseline fluorescence F_0_ being defined as the mean pixel intensity of the 2 second time window preceding the optogenetic stimulus.

### Statistics

No statistical methods were used to predetermine sample size. The sample size was determined based on the signal strength and variability in pilot experiments. In the larva, data were collected from all the tested samples according to the screening procedure detailed in the section *Calcium imaging in the larva*. In the adult, data were collected from all the tested samples. During each experiment the indicated light stimulus was repeated 5 times (Figures 1 and 3) or 6 times (Figure 2). In the trace-plots (for example Figure 1B) the faint traces indicate the median of a single experiment. The bold trace is the mean of all experiments. In the paired-data plots (for example Figure 1C) each point is the median of a single experiment: ‘Before’ is the median ΔF/F of the second preceding the optogenetic stimulus. ‘During’ is the median ΔF/F for the last 1 second of the stimulation period in the larva (Figures 1 and 3). In adult ‘During’ is the median ΔF/F of the 10 seconds of the stimulus.

To test whether the optogenetic stimulation leads to an increase in ΔF/F the data between ‘Before’ and ‘During’ was compared. To test for normality the Lilliefors test (Python function: statsmodels.stats.diagnostic.lilliefors) was used with standard settings. To test for equality of variances Levene’s test (Python function: scipy.stats.levene) was used with standard settings. To test for differences in ΔF/F before and during the stimulus, the dependent t-test for paired samples (scipy.stats.ttest_rel) was used with standard settings. Multiple test correction was done using the Holm-Bonferroni method.

## Data availability

*Drosophila* transgenic trains used in this study are available either from the Bloomington Drosophila Stock Center or upon request from the authors. Custom scripts to plot all figures and run all statistical tests in this manuscript from the included dF/F traces are available through the following repository: https://gitlab.com/davidtadres/leaky_expression_publication.

## Acknowledgments

We thank V. Jayaraman, B. Dickson as well as members from Simpson and Kim labs at UCSB for discussion. We thank C. Ribeiro to support the work of IT. We are grateful to Todd Laverty,the Janelia fly and molecular biology cores for assistance with the generation of new fly stocks. This work was supported by the National Institute for Health (RO1-NS113048-01) (DT and ML), the University of California, Santa Barbara (startup funds) (DT and ML), the Janelia Research Campus Visitor Program (DT and ML) and the Howard Hughes Medical Institute (HS and DS).

---

**Supplementary Figure 1:**
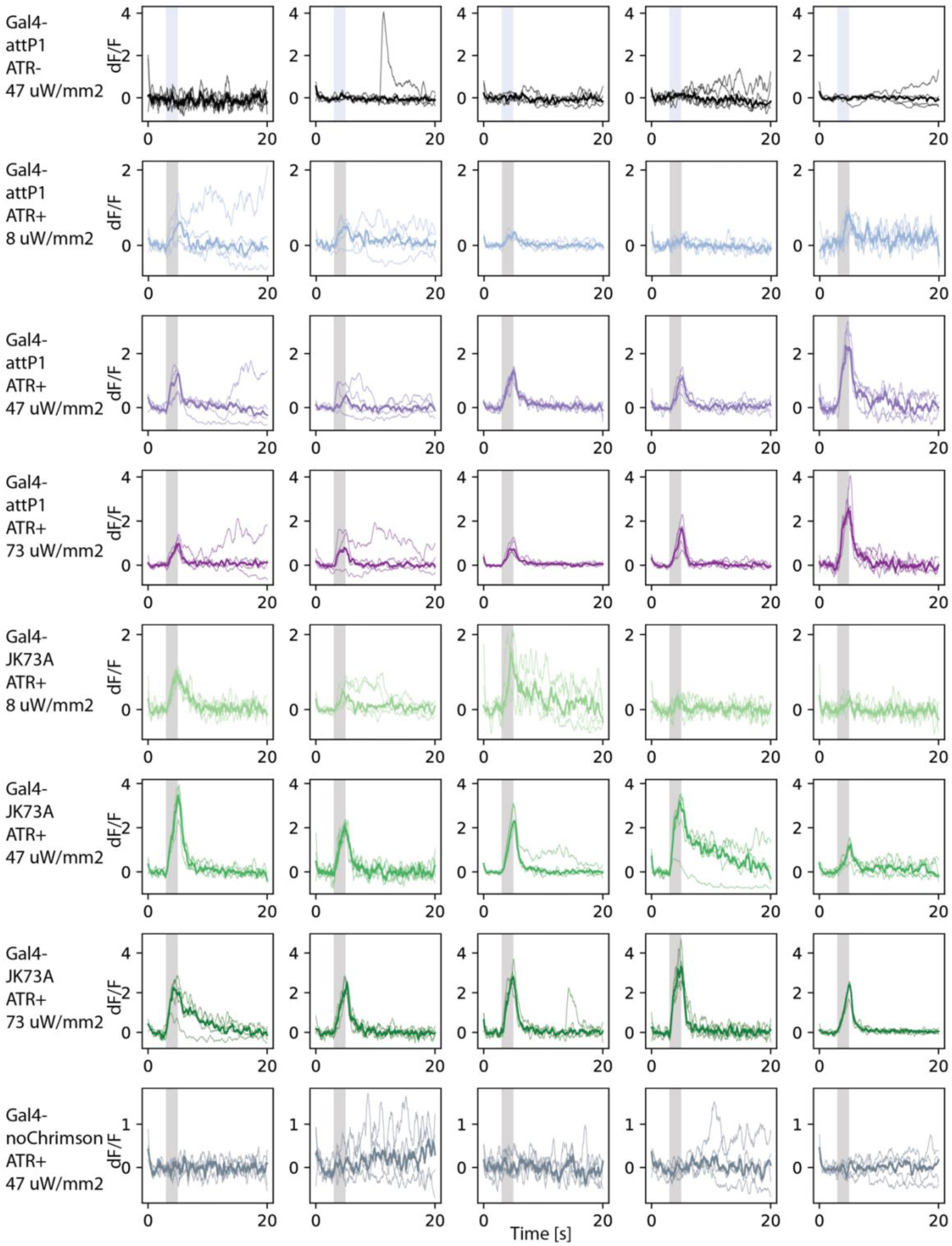
Overview of raw data shown in Figure 1. Each row contains the individual animals of the genotype indicated on the left. Each box shows the ΔF/F over time for each of the 5 repeats with the thinner lines in the background. The median ΔF/F is shown using the thick line.

**Supplementary Figure 2:**
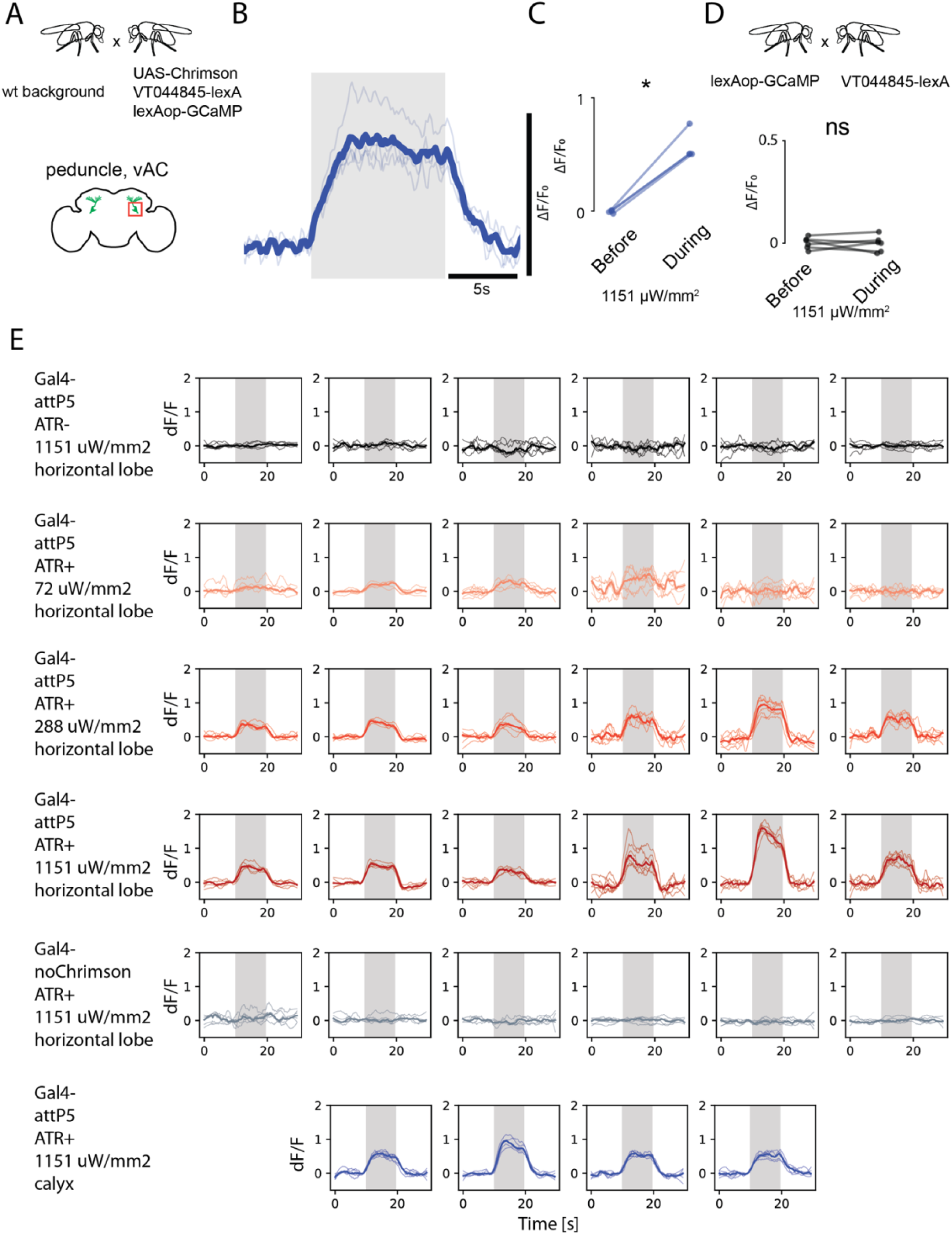
Leaky expression of UAS-Chrimson in the adult fly brain. **(A)** Calcium imaging was performed in the calyx and peduncle. **(B)** Calcium transients of adult flies grown on food with ATR (red). Grey shade indicates 660 nm stimulus with power of 1151 uW/mm2. The thick line represents the median of the calcium signals computed over the individual trials (thin lines). **(C)** Flies with the UAS-Chrimson transgene in su(Hw)attP5 have a significant increase in ΔF/F upon stimulation (n=4 animals, paired t-test, p<0.05). **(D)** Data from adult flies with no Gal4 driver (n=6 animals, paired t-test, p>0.05). **(E)** Overview of raw data shown in panels B and C, as well as in Figure 2. Each row contains the individual animals of the genotype and imaging location indicated on the left. Each box shows the ΔF/F over time for each of the 6 or 4 (horizontal lobe and calyx, respectively) repeats with the thinner lines in the background. The median ΔF/F is shown using the thick line.

**Supplementary Figure 3:**
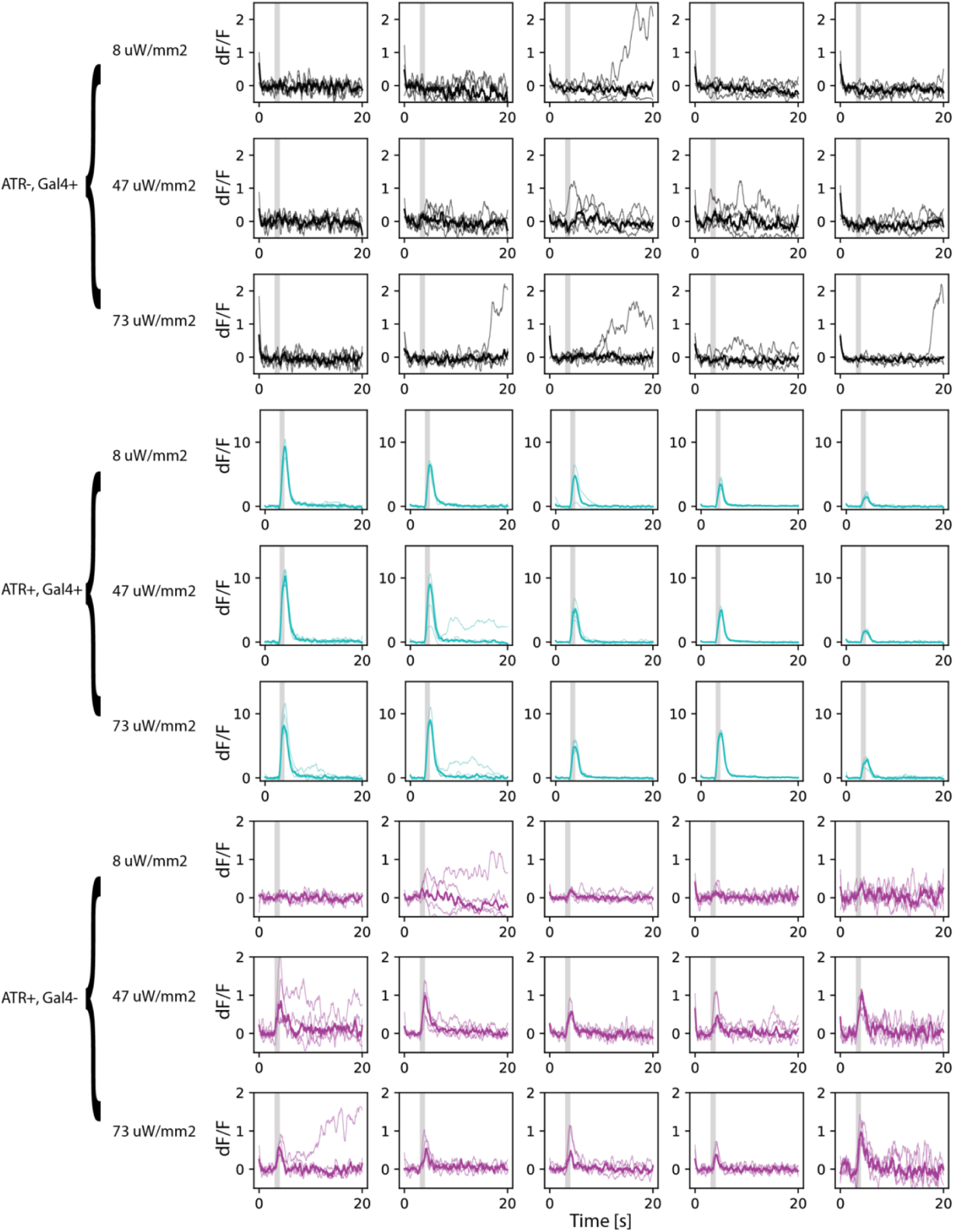
Overview of raw data shown in Figure 3. Each row contains the individual animals of the genotype and light intensity indicated on the left. Each box shows the ΔF/F over time for each of the 5 repeats with the thinner lines in the background. The median ΔF/F is shown using the thick line.

## References

Aso, Y., Hattori, D., Yu, Y., Johnston, R.M., Iyer, N.A., Ngo, T.-T., Dionne, H., Abbott, L., Axel, R., Tanimoto, H., Rubin, G.M., 2014. The neuronal architecture of the mushroom body provides a logic for associative learning. eLife 3, e04577. https://doi.org/10.7554/eLife.04577

Borden, P.M., Zhang, P., Shivange, A.V., Marvin, J.S., Cichon, J., Dan, C., Podgorski, K., Figueiredo, A., Novak, O., Tanimoto, M., Shigetomi, E., Lobas, M.A., Kim, H., Zhu, P.K., Zhang, Y., Zheng, W.S., Fan, C., Wang, G., Xiang, B., Gan, L., Zhang, G.-X., Guo, K., Lin, L., Cai, Y., Yee, A.G., Aggarwal, A., Ford, C.P., Rees, D.C., Dietrich, D., Khakh, B.S., Dittman, J.S., Gan, W.-B., Koyama, M., Jayaraman, V., Cheer, J.F., Lester, H.A., Zhu, J.J., Looger, L.L., 2020. A fast genetically encoded fluorescent sensor for faithful in vivo acetylcholine detection in mice, fish, worms and flies. https://doi.org/10.1101/2020.02.07.939504

Borst, A., Helmstaedter, M., 2015. Common circuit design in fly and mammalian motion vision. Nat Neurosci 18, 1067–1076. https://doi.org/10.1038/nn.4050

Brand, A.H., Perrimon, N., 1993. Targeted gene expression as a means of altering cell fates and generating dominant phenotypes. Development 118, 401–415.

Buhmann, J., Sheridan, A., Malin-Mayor, C., Schlegel, P., Gerhard, S., Kazimiers, T., Krause, R., Nguyen, T.M., Heinrich, L., Lee, W.-C.A., Wilson, R., Saalfeld, S., Jefferis, G.S.X.E., Bock, D.D., Turaga, S.C., Cook, M., Funke, J., 2021. Automatic detection of synaptic partners in a whole-brain Drosophila electron microscopy data set. Nat Methods 18, 771–774. https://doi.org/10.1038/s41592-021-01183-7

Carreira-Rosario, A., Zarin, A.A., Clark, M.Q., Manning, L., Fetter, R.D., Cardona, A., Doe, C.Q., 2018. MDN brain descending neurons coordinately activate backward and inhibit forward locomotion. eLife 7, e38554. https://doi.org/10.7554/eLife.38554

Chamberland, S., Yang, H.H., Pan, M.M., Evans, S.W., Guan, S., Chavarha, M., Yang, Y., Salesse, C., Wu, H., Wu, J.C., Clandinin, T.R., Toth, K., Lin, M.Z., St-Pierre, F., 2017. Fast two-photon imaging of subcellular voltage dynamics in neuronal tissue with genetically encoded indicators. eLife 6, e25690. https://doi.org/10.7554/eLife.25690

Chen, C., Agrawal, S., Mark, B., Mamiya, A., Sustar, A., Phelps, J.S., Lee, W.-C.A., Dickson, B.J., Card, G.M., Tuthill, J.C., 2021. Functional architecture of neural circuits for leg proprioception in Drosophila. Current Biology. https://doi.org/10.1016/j.cub.2021.09.035

Chen, T.-W., Wardill, T.J., Sun, Y., Pulver, S.R., Renninger, S.L., Baohan, A., Schreiter, E.R., Kerr, R.A., Orger, M.B., Jayaraman, V., Looger, L.L., Svoboda, K., Kim, D.S., 2013. Ultrasensitive fluorescent proteins for imaging neuronal activity. Nature 499, 295–300. https://doi.org/10.1038/nature12354

De Oliveira, G.P.R., Rodriguez-Amaya, D.B., 2007. Processed and Prepared Corn Products As Sources of Lutein and Zeaxanthin: Compositional Variation in the Food Chain. Journal of Food Science 72, S079–S085. https://doi.org/10.1111/j.1750-3841.2006.00235.x

Dorkenwald, S., McKellar, C.E., Macrina, T., Kemnitz, N., Lee, K., Lu, R., Wu, J., Popovych, S., Mitchell, E., Nehoran, B., Jia, Z., Bae, J.A., Mu, S., Ih, D., Castro, M., Ogedengbe, O., Halageri, A., Kuehner, K., Sterling, A.R., Ashwood, Z., Zung, J., Brittain, D., Collman, F., Schneider-Mizell, C., Jordan, C., Silversmith, W., Baker, C., Deutsch, D., Encarnacion-Rivera, L., Kumar, S., Burke, A., Bland, D., Gager, J., Hebditch, J., Koolman, S., Moore, M., Morejohn, S., Silverman, B., Willie, K., Willie, R., Yu, S., Murthy, M., Seung, H.S., 2022. FlyWire: online community for whole-brain connectomics. Nat Methods 19, 119–128. https://doi.org/10.1038/s41592-021-01330-0

Eichler, K., Li, F., Litwin-Kumar, A., Park, Y., Andrade, I., Schneider-Mizell, C.M., Saumweber, T., Huser, A., Eschbach, C., Gerber, B., Fetter, R.D., Truman, J.W., Priebe, C.E., Abbott, L.F., Thum, A.S., Zlatic, M., Cardona, A., 2017. The complete connectome of a learning and memory centre in an insect brain. Nature 548, 175–182. https://doi.org/10.1038/nature23455

Eschbach, C., Fushiki, A., Winding, M., Afonso, B., Andrade, I.V., Cocanougher, B.T., Eichler, K., Gepner, R., Si, G., Valdes-Aleman, J., Fetter, R.D., Gershow, M., Jefferis, G.S., Samuel, A.D., Truman, J.W., Cardona, A., Zlatic, M., 2021. Circuits for integrating learned and innate valences in the insect brain. eLife 10, e62567. https://doi.org/10.7554/eLife.62567

Eschbach, C., Zlatic, M., 2020. Useful road maps: studying Drosophila larva’s central nervous system with the help of connectomics. Current Opinion in Neurobiology, Whole-brain interactions between neural circuits 65, 129–137. https://doi.org/10.1016/j.conb.2020.09.008

Franconville, R., Beron, C., Jayaraman, V., 2018. Building a functional connectome of the Drosophila central complex. eLife 7, e37017. https://doi.org/10.7554/eLife.37017

Gowda, S.B.M., Salim, S., Mohammad, F., 2021. Anatomy and Neural Pathways Modulating Distinct Locomotor Behaviors in Drosophila Larva. Biology 10, 90. https://doi.org/10.3390/biology10020090

Hayashi, S., Ito, K., Sado, Y., Taniguchi, M., Akimoto, A., Takeuchi, H., Aigaki, T., Matsuzaki, F., Nakagoshi, H., Tanimura, T., Ueda, R., Uemura, T., Yoshihara, M., Goto, S., 2002. GETDB, a database compiling expression patterns and molecular locations of a collection of gal4 enhancer traps. genesis 34, 58–61. https://doi.org/10.1002/gene.10137

Hulse, B.K., Haberkern, H., Franconville, R., Turner-Evans, D.B., Takemura, S., Wolff, T., Noorman, M., Dreher, M., Dan, C., Parekh, R., Hermundstad, A.M., Rubin, G.M., Jayaraman, V., 2021. A connectome of the Drosophila central complex reveals network motifs suitable for flexible navigation and context-dependent action selection. eLife 10, e66039. https://doi.org/10.7554/eLife.66039

Jenett, A., Rubin, G.M., Ngo, T.-T.B., Shepherd, D., Murphy, C., Dionne, H., Pfeiffer, B.D., Cavallaro, A., Hall, D., Jeter, J., Iyer, N., Fetter, D., Hausenfluck, J.H., Peng, H., Trautman, E.T., Svirskas, R.R., Myers, E.W., Iwinski, Z.R., Aso, Y., DePasquale, G.M., Enos, A., Hulamm, P., Lam, S.C.B., Li, H.-H., Laverty, T.R., Long, F., Qu, L., Murphy, S.D., Rokicki, K., Safford, T., Shaw, K., Simpson, J.H., Sowell, A., Tae, S., Yu, Y., Zugates, C.T., 2012. A GAL4-Driver Line Resource for Drosophila Neurobiology. Cell Reports 2, 991–1001. https://doi.org/10.1016/j.celrep.2012.09.011

Jia, H., Rochefort, N.L., Chen, X., Konnerth, A., 2011. In vivo two-photon imaging of sensory-evoked dendritic calcium signals in cortical neurons. Nature Protocols 6, 28–35. https://doi.org/10.1038/nprot.2010.169

Jing, M., Li, Yuexuan, Zeng, J., Huang, P., Skirzewski, M., Kljakic, O., Peng, W., Qian, T., Tan, K., Zou, J., Trinh, S., Wu, R., Zhang, S., Pan, S., Hires, S.A., Xu, M., Li, H., Saksida, L.M., Prado, V.F., Bussey, T.J., Prado, M.A.M., Chen, L., Cheng, H., Li, Yulong, 2020. An optimized acetylcholine sensor for monitoring in vivo cholinergic activity. Nat Methods 17, 1139–1146. https://doi.org/10.1038/s41592-020-0953-2

Keller, A., Sweeney, S.T., Zars, T., O’Kane, C.J., Heisenberg, M., 2002. Targeted expression of tetanus neurotoxin interferes with behavioral responses to sensory input in Drosophila. Journal of Neurobiology 50, 221–233. https://doi.org/10.1002/neu.10029

Klapoetke, N.C., Murata, Y., Kim, S.S., Pulver, S.R., Birdsey-Benson, A., Cho, Y.K., Morimoto, T.K., Chuong, A.S., Carpenter, E.J., Tian, Z., Wang, J., Xie, Y., Yan, Z., Zhang, Y., Chow, B.Y., Surek, B., Melkonian, M., Jayaraman, V., Constantine-Paton, M., Wong, G.K.-S., Boyden, E.S., 2014. Independent optical excitation of distinct neural populations. Nature Methods 11, 338–346. https://doi.org/10.1038/nmeth.2836

Knapp, J.-M., Chung, P., Simpson, J.H., 2015. Generating Customized Transgene Landing Sites and Multi-Transgene Arrays in Drosophila Using phiC31 Integrase. Genetics 199, 919–934. https://doi.org/10.1534/genetics.114.173187

Lai, S.-L., Lee, T., 2006. Genetic mosaic with dual binary transcriptional systems in Drosophila. Nat Neurosci 9, 703–709. https://doi.org/10.1038/nn1681

Li, H.-H., Kroll, J.R., Lennox, S.M., Ogundeyi, O., Jeter, J., Depasquale, G., Truman, J.W., 2014. A GAL4 Driver Resource for Developmental and Behavioral Studies on the Larval CNS of Drosophila. Cell Reports 8, 897–908. https://doi.org/10.1016/j.celrep.2014.06.065

Lillvis, J.L., Otsuna, H., Ding, X., Pisarev, I., Kawase, T., Colonell, J., Rokicki, K., Goina, C., Gao, R., Hu, A., Wang, K., Bogovic, J., Milkie, D.E., Meienberg, L., Boyden, E.S., Saalfeld, S., Tillberg, P.W., Dickson, B.J., 2021. Rapid reconstruction of neural circuits using tissue expansion and lattice light sheet microscopy. https://doi.org/10.1101/2021.11.14.468535

Markstein, M., Pitsouli, C., Villalta, C., Celniker, S.E., Perrimon, N., 2008. Exploiting position effects and the gypsy retrovirus insulator to engineer precisely expressed transgenes. Nat Genet 40, 476–483. https://doi.org/10.1038/ng.101

Meinertzhagen, I.A., 2018. Of what use is connectomics? A personal perspective on the Drosophila connectome. Journal of Experimental Biology 221, jeb164954. https://doi.org/10.1242/jeb.164954

Oberhauser, V., Voolstra, O., Bangert, A., von Lintig, J., Vogt, K., 2008. NinaB combines carotenoid oxygenase and retinoid isomerase activity in a single polypeptide. Proceedings of the National Academy of Sciences 105, 19000–19005. https://doi.org/10.1073/pnas.0807805105

Pfeiffer, B.D., Ngo, T.-T.B., Hibbard, K.L., Murphy, C., Jenett, A., Truman, J.W., Rubin, G.M., 2010. Refinement of Tools for Targeted Gene Expression in Drosophila. Genetics 186, 735–755. https://doi.org/10.1534/genetics.110.119917

Pnevmatikakis, E.A., Giovannucci, A., 2017. NoRMCorre: An online algorithm for piecewise rigid motion correction of calcium imaging data. Journal of Neuroscience Methods 291, 83–94. https://doi.org/10.1016/j.jneumeth.2017.07.031

Potter, C.J., Tasic, B., Russler, E.V., Liang, L., Luo, L., 2010. The Q System: A Repressible Binary System for Transgene Expression, Lineage Tracing, and Mosaic Analysis. Cell 141, 536–548. https://doi.org/10.1016/j.cell.2010.02.025

Qiao, H.-H., Wang, F., Xu, R.-G., Sun, J., Zhu, R., Mao, D., Ren, X., Wang, X., Jia, Y., Peng, P., Shen, D., Liu, L.-P., Chang, Z., Wang, G., Li, S., Ji, J.-Y., Liu, Q., Ni, J.-Q., 2018. An efficient and multiple target transgenic RNAi technique with low toxicity in Drosophila. Nat Commun 9, 4160. https://doi.org/10.1038/s41467-018-06537-y

Randel, N., Shahidi, R., Verasztó, C., Bezares-Calderón, L.A., Schmidt, S., Jékely, G., 2015. Inter-individual stereotypy of the Platynereis larval visual connectome. eLife 4, e08069. https://doi.org/10.7554/eLife.08069

Scheffer, L.K., Xu, C.S., Januszewski, M., Lu, Z., Takemura, Shin-ya, Hayworth, K.J., Huang, G.B., Shinomiya, K., Maitlin-Shepard, J., Berg, S., Clements, J., Hubbard, P.M., Katz, W.T., Umayam, L., Zhao, T., Ackerman, D., Blakely, T., Bogovic, J., Dolafi, T., Kainmueller, D., Kawase, T., Khairy, K.A., Leavitt, L., Li, P.H., Lindsey, L., Neubarth, N., Olbris, D.J., Otsuna, H., Trautman, E.T., Ito, M., Bates, A.S., Goldammer, J., Wolff, T., Svirskas, R., Schlegel, P., Neace, E., Knecht, C.J., Alvarado, C.X., Bailey, D.A., Ballinger, S., Borycz, J.A., Canino, B.S., Cheatham, N., Cook, M., Dreher, M., Duclos, O., Eubanks, B., Fairbanks, K., Finley, S., Forknall, N., Francis, A., Hopkins, G.P., Joyce, E.M., Kim, S., Kirk, N.A., Kovalyak, J., Lauchie, S.A., Lohff, A., Maldonado, C., Manley, E.A., McLin, S., Mooney, C., Ndama, M., Ogundeyi, O., Okeoma, N., Ordish, C., Padilla, N., Patrick, C.M., Paterson, T., Phillips, E.E., Phillips, E.M., Rampally, N., Ribeiro, C., Robertson, M.K., Rymer, J.T., Ryan, S.M., Sammons, M., Scott, A.K., Scott, A.L., Shinomiya, A., Smith, C., Smith, K., Smith, N.L., Sobeski, M.A., Suleiman, A., Swift, J., Takemura, Satoko, Talebi, I., Tarnogorska, D., Tenshaw, E., Tokhi, T., Walsh, J.J., Yang, T., Horne, J.A., Li, F., Parekh, R., Rivlin, P.K., Jayaraman, V., Costa, M., Jefferis, G.S., Ito, K., Saalfeld, S., George, R., Meinertzhagen, I.A., Rubin, G.M., Hess, H.F., Jain, V., Plaza, S.M., 2020. A connectome and analysis of the adult Drosophila central brain. eLife 9, e57443. https://doi.org/10.7554/eLife.57443

Schlegel, P., Costa, M., Jefferis, G.S., 2017. Learning from connectomics on the fly. Current Opinion in Insect Science, Neuroscience * Pheromones 24, 96–105. https://doi.org/10.1016/j.cois.2017.09.011

Schneider-Mizell, C.M., Gerhard, S., Longair, M., Kazimiers, T., Li, F., Zwart, M.F., Champion, A., Midgley, F.M., Fetter, R.D., Saalfeld, S., Cardona, A., 2016. Quantitative neuroanatomy for connectomics in Drosophila. eLife 5, e12059. https://doi.org/10.7554/eLife.12059

Scholz, H., Ramond, J., Singh, C.M., Heberlein, U., 2000. Functional Ethanol Tolerance in Drosophila. Neuron 28, 261–271. https://doi.org/10.1016/S0896-6273(00)00101-X

Sen, R., Wu, M., Branson, K., Robie, A., Rubin, G.M., Dickson, B.J., 2017. Moonwalker Descending Neurons Mediate Visually Evoked Retreat in Drosophila. Current Biology 27, 766–771. https://doi.org/10.1016/j.cub.2017.02.008

Simpson, J.H., 2009. Chapter 3 Mapping and Manipulating Neural Circuits in the Fly Brain, in: Advances in Genetics, Genetic Dissection of Neural Circuits and Behavior. Academic Press, pp. 79–143. https://doi.org/10.1016/S0065-2660(09)65003-3

Simpson, J.H., Looger, L.L., 2018. Functional Imaging and Optogenetics in Drosophila. Genetics 208, 1291–1309. https://doi.org/10.1534/genetics.117.300228

Takagi, S., Cocanougher, B.T., Niki, S., Miyamoto, D., Kohsaka, H., Kazama, H., Fetter, R.D., Truman, J.W., Zlatic, M., Cardona, A., Nose, A., 2017. Divergent Connectivity of Homologous Command-like Neurons Mediates Segment-Specific Touch Responses in Drosophila. Neuron 96, 1373-1387.e6. https://doi.org/10.1016/j.neuron.2017.10.030

Takemura, Shin-ya, Bharioke, A., Lu, Z., Nern, A., Vitaladevuni, S., Rivlin, P.K., Katz, W.T., Olbris, D.J., Plaza, S.M., Winston, P., Zhao, T., Horne, J.A., Fetter, R.D., Takemura, Satoko, Blazek, K., Chang, L.-A., Ogundeyi, O., Saunders, M.A., Shapiro, V., Sigmund, C., Rubin, G.M., Scheffer, L.K., Meinertzhagen, I.A., Chklovskii, D.B., 2013. A visual motion detection circuit suggested by Drosophila connectomics. Nature 500, 175–181. https://doi.org/10.1038/nature12450

Tang, X., Roessingh, S., Hayley, S.E., Chu, M.L., Tanaka, N.K., Wolfgang, W., Song, S., Stanewsky, R., Hamada, F.N., 2017. The role of PDF neurons in setting the preferred temperature before dawn in Drosophila. eLife 6, e23206. https://doi.org/10.7554/eLife.23206

Tastekin, I., Khandelwal, A., Tadres, D., Fessner, N.D., Truman, J.W., Zlatic, M., Cardona, A., Louis, M., 2018. Sensorimotor pathway controlling stopping behavior during chemotaxis in the Drosophila melanogaster larva. eLife 7, e38740. https://doi.org/10.7554/eLife.38740

Tirian, L., Dickson, B.J., 2017. The VT GAL4, LexA, and split-GAL4 driver line collections for targeted expression in the Drosophila nervous system. https://doi.org/10.1101/198648

Vogt, K., Aso, Y., Hige, T., Knapek, S., Ichinose, T., Friedrich, A.B., Turner, G.C., Rubin, G.M., Tanimoto, H., 2016. Direct neural pathways convey distinct visual information to Drosophila mushroom bodies. eLife 5, e14009. https://doi.org/10.7554/eLife.14009

Wang, K., Wang, F., Forknall, N., Yang, T., Patrick, C., Parekh, R., Dickson, B.J., 2021. Neural circuit mechanisms of sexual receptivity in Drosophila females. Nature 589, 577–581. https://doi.org/10.1038/s41586-020-2972-7

Wanner, A.A., Genoud, C., Masudi, T., Siksou, L., Friedrich, R.W., 2016. Dense EM-based reconstruction of the interglomerular projectome in the zebrafish olfactory bulb. Nat Neurosci 19, 816–825. https://doi.org/10.1038/nn.4290

Yoshino, J., Morikawa, R.K., Hasegawa, E., Emoto, K., 2017. Neural Circuitry that Evokes Escape Behavior upon Activation of Nociceptive Sensory Neurons in Drosophila Larvae. Current Biology 27, 2499-2504.e3. https://doi.org/10.1016/j.cub.2017.06.068

Zheng, Z., Lauritzen, J.S., Perlman, E., Robinson, C.G., Nichols, M., Milkie, D., Torrens, O., Price, J., Fisher, C.B., Sharifi, N., Calle-Schuler, S.A., Kmecova, L., Ali, I.J., Karsh, B., Trautman, E.T., Bogovic, J.A., Hanslovsky, P., Jefferis, G.S.X.E., Kazhdan, M., Khairy, K., Saalfeld, S., Fetter, R.D., Bock, D.D., 2018. A Complete Electron Microscopy Volume of the Brain of Adult Drosophila melanogaster. Cell 174, 730-743.e22. https://doi.org/10.1016/j.cell.2018.06.019

